# G-quadruplex prediction in *E.coli* genome reveals a conserved putative G-quadruplex-Hairpin-Duplex switch

**DOI:** 10.1101/032615

**Authors:** Osman Doluca, Oktay I Kaplan, Burak Berber, Nezih Hekim

## Abstract

Many studies show that short non-coding sequences are widely conserved among regulatory elements. More and more conserved sequences are being discovered since the development next generation sequencing technology. A common approach to identify conserved sequences with regulatory roles rely on the topological change such as hairpin formation on the DNA or RNA level. However, G-quadruplexes, a non-cannonical nucleic acid topology with little established biological role, is rarely considered for conserved regulatory element discovery. Here we present the use of G-quadruplex prediction algorithm to identify putative G-quadruplex-forming and conserved elements in E.coli genome. Phylogenetic analysis of 52 G-quadruplex forming sequences revealed two conserved G-quadruplex motifs with potential regulatory role.

## Introduction

Short, non-coding and recurring sequences, referred as sequence motifs, often indicate a protein binding site or a site for a structural modification that eventually leads to the alteration of interaction with enzymes. The latter, also referred as structural motifs may be detected based on mathematical prediction of putative alternate topologies, rather than common B-form (Hatfield and Benham 2002; Perez-Martin and de Lorenzo 1997). Especially since the development of next generation sequencing technologies we have an unprecedented amount of sequence data waiting to be scanned for such topologies. While the most structural motif prediction algorithms rely on Watson-crick base-pairing and associated topologies, only a few tools are available focusing on scanning for alternate nucleic acid structures.

The G-quadruplexes (GQs) is a large group of evolutionarily conserved higher order structures present in both lower (bacteria)(Rawal et al. 2006) and higher organisms (worms and humans) (Murat and Balasubramanian 2014; Patel, Phan, and Kuryavyi 2007). G-quadruplexes are formed by guanine-rich sequences and defined in general terms as structures formed by a core of stacked G-tetrads, plenary aligned guanines held by Hoogsteen H-bonding, and connected by loops arising from intervening mixed sequences that are not usually involved in tetrads themselves(Gellert, Lipsett, and Davies 1962). The combination of the number of stacked tetrads, the polarity of the strands, the location and length of the loops as well as *syn* vs *anti* conformation of the guanine bases leads to a vast variety of G-quadruplex topologies.

The GQs were first discovered *in vitro* and then in eukaryotes, in telomeres and more recently in non-telomeric promoters and introns (Burge et al. 2006). Biological significance of GQs recently started to emerge long after their *in vitro* discovery. For instance, telomeric GQs that are formed by repeating dTAAGGG sequences is responsible for maintaining telomere length and telomere stability while G-rich sequences that are formed in promoters of oncogenes such as KRAS and c-MYC, are associated with gene regulation (Simonsson, Pecinka, and Kubista 1998). Because GQs may be recognized by transcription factors, the transcription of these genes can be regulated by G-quadruplex binding molecules.

While several studies have focused to identify GQ motifs and their roles in eukaryotes, there hasn’t been near enough study on GQs in prokaryotes. A bioinformatics study on GQs in *E. coli* has indicated a regulatory role for GQs found in regulatory regions (−200 to −1 bp upstream from transcription start site, TSS) (Rawal et al. 2006). Here, we have applied a more restrict algorithm to identify 52 highly putative G-quadruplex forming (HPGQ) motifs in *E.coli*. Phylogenetic grouping of these sequences revealed two groups of highly conserved HPGQs with 16 and 7 members. To our surprise all members of both groups were located only in intergenic regions and especially close to the 3’UTR and mostly between operons, indicating a regulatory role. Due to their highly conserved sequence HPG1 and HPG2 are expected to form a unique topology.

## Results

### Identification of highly putative G-quadruplexes

A G-quadruplex actually may refer to a large group of topologies with a common feature of G-tetrad, a non-canonical assembly of four guanines on a plane. Because the stacks of G-tetrads are required to stabilize the structure, four repeats, or tracts, of guanines are necessary in any sequence to form an intramolecular GQ. While the length of these repeats has the biggest impact on the stability, so do the sequences linking the guanine tracts (G-tracts). In the light of this information, a generally accepted algorithm of G*_S_*N*_L1_*G*_S_*N*_L2_*G*_S_*N*_L3_*G*_S_* is used to determine the putative G-quadruplex forming sequences, where *S* refers to the length of the GQ stem, *L1-3* defines the lengths of the loops and are independent from each other. While the effect of loop lengths on the ability of GQ formation is highly debatable, the repeat length, or stem length, is expectedly highly influential on the GQ formation due to cumulative *π*-*π* interactions. In most cases a repeat length of 3 is considered a stringent rule for the formation of GQ *in vivo*. In order to capture only highly putative G-quadruplexes we restricted *S* to be above or equal to 3 while *L1-7* are set to be between 1 and 7. A whole genome scan using this algorithm yielded only 52 sequences in *E.coli* genome (K12 MG1655 strain). These highly putative G-quadruplexes are abbreviated as HPGQs and listed in Table 1S. In cases of sequences with overlapping patterns due to presence of more than four guanine tracts, we selected the largest patterns to include all G-tracts. It is important to note that such patterns may adopt multiple topologies depending on the G-tracts that take part in the G-tetrad formations.

### Phylogenetic classification of HPGQs

We have previously mentioned that a G-quadruplex may have a very large variety of topologies. When investigating the role of a GQ, this topological variety should be taken into account because any protein-GQ interaction would strongly be dependent on the topology of the GQ. Since the topology is strongly controlled by the sequence, it is possible to relate the sequence of a GQ to its function. This is especially possible when other factors are also identical (*ie*. intracellular cation concentration). Thus, in order to classify the GQs, we hypothesized of using a phylogenetic analysis to group the HPGQs of *E.coli*. The sequences of HPGQs were aligned using Clustal Omega web tool (Goujon et al. 2010; Thompson, Gibson, and Higgins 2002). With a phylogenetic distance threshold of 0.047 the sequences were grouped into two distinct HPGQ groups with 16 and 7 members, HPG1 and HPG2, respectively (Figure 1). The threshold of 0.047 as the ratio of the number of permitted substitutions to the length of alignment, allowed up to a single mismatch or insertion/deletion in the sequences. The rest of HPGQs had unique sequences and high phylogenetic distance between each other and left ungrouped. Our research continued on the two groups that are identified at this point by this alignment as HPG1 and HPG2

**Figure 1.**
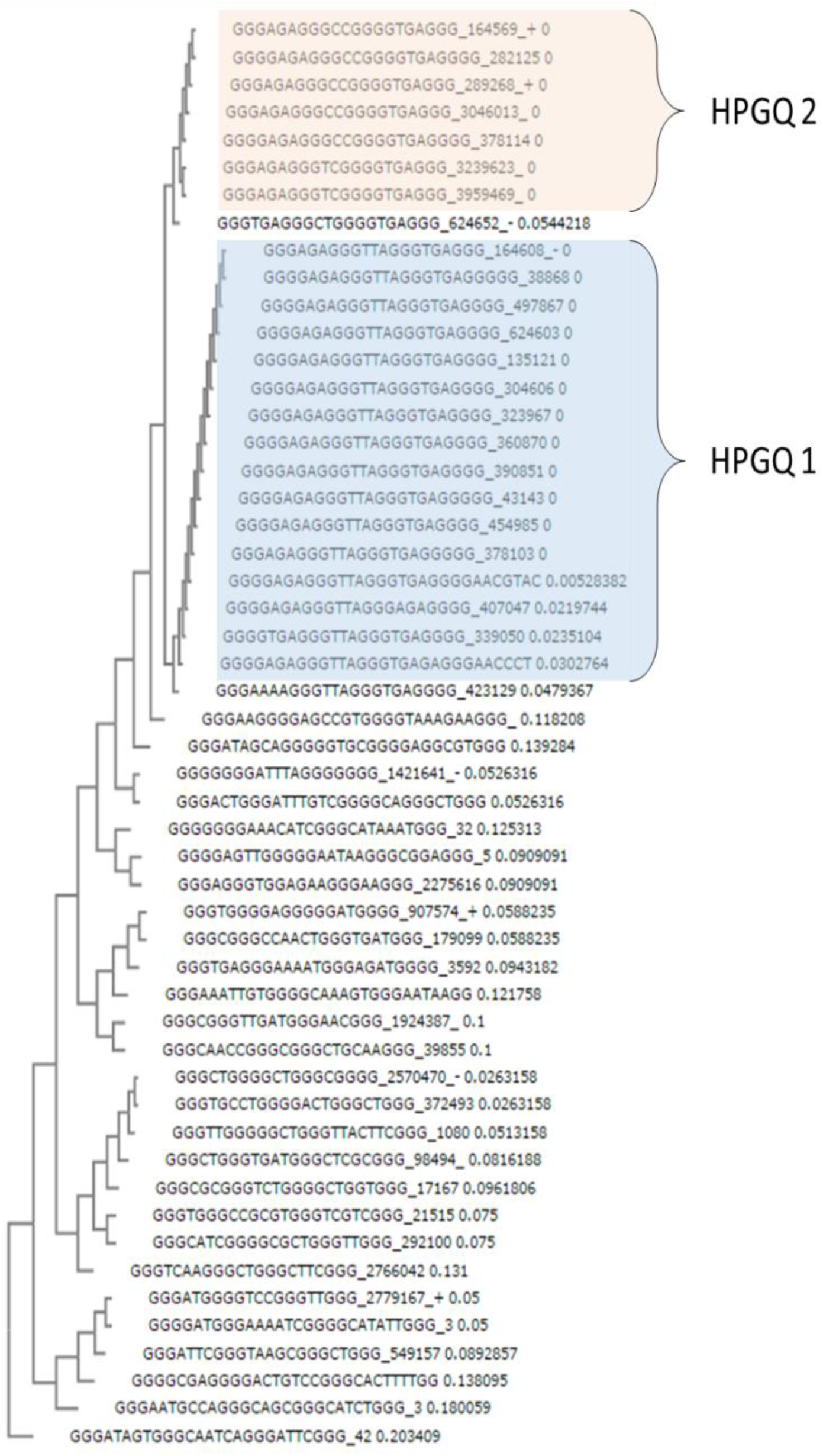
*Phylogenetic tree of E.coli HPGQs*.

Interestingly the alignment revealed that 13 of HPG1 and another 5 of HPG2 had the identical conserved HPGQ sequences within each group; dGGGAGAGGGTTAGGGTGAGGG (cHPG1) and dGGGAGAGGGCCGGGGTGAGGG (cHPG2), respectively. Moreover, cHPG1, cHPG2, showed striking similarity with only difference in the second loop (dTTA vs dCC). Because the difference is only on the middle loop, it may have a distinguishable topological difference only on a single face of the G-quadruplex (*ie*. top or bottom), thus may have a similar functionality.

### Genomic distribution of HPGQs in *E.coli*

We identified the location of HPGQ motifs of HPG1 and HPG2 according to the nearest open reading frames (ORFs) and operons of *E.coli* genome. Surprisingly, all motifs are found to be located at sites flanking the ORFs and none within. Their mid-point relative to the nearest genes, the directions of the genes and the strand of the motifs are listed in Table 1.

**Table 1.**
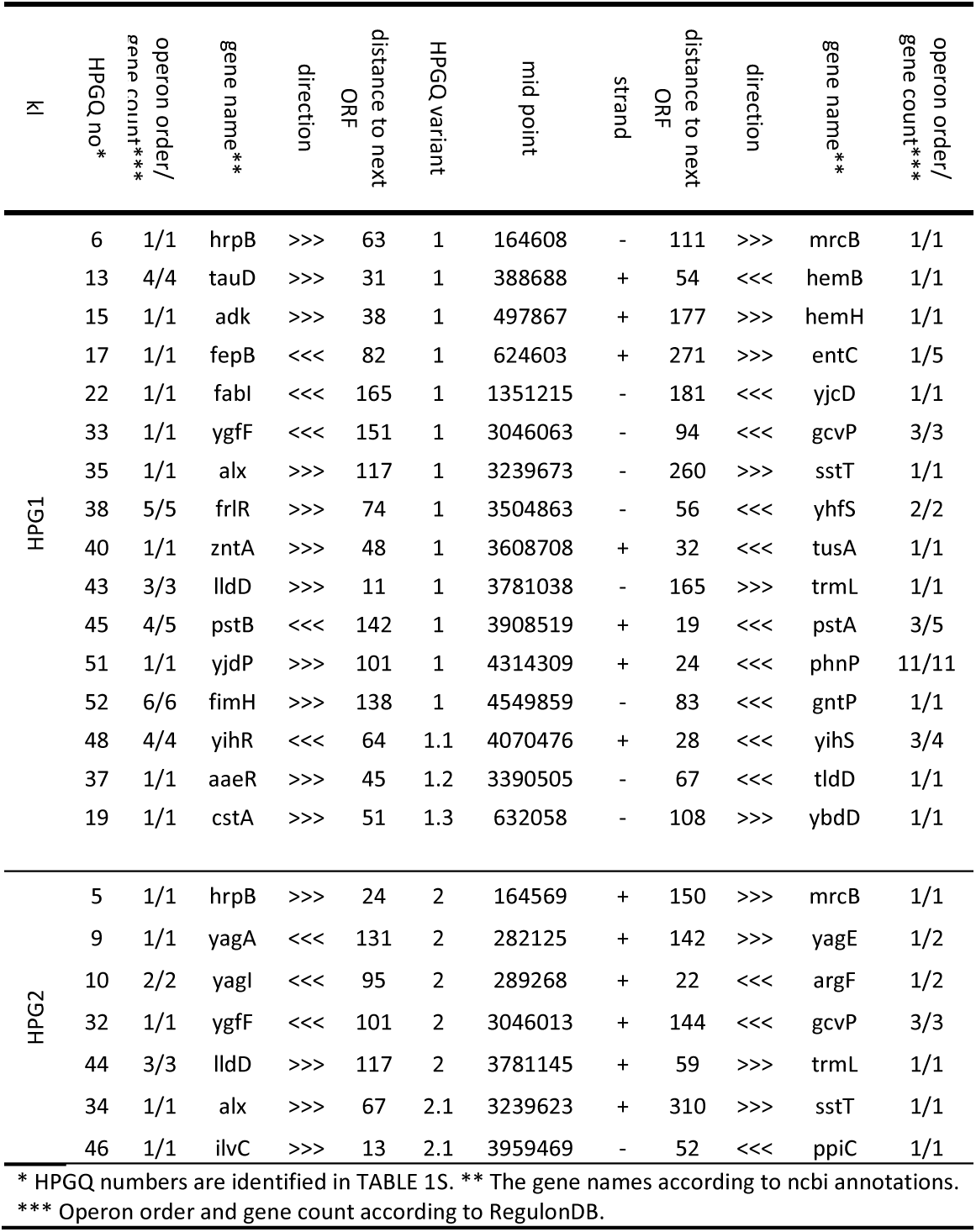
*List of HPGQs positions and relative positions of the related genes*.

HPGQs are identified according to directionality of the neighbouring genes as between tandem, convergent or divergent gene pairs. The most of the HPGQs are found to be between tandem gene pairs and often positioned closer to the end than the beginning of the ORFs. Figure 2 shows the numbers of HPG1s (A) and HPG2s (B) found between tandem, convergent or divergent gene pairs.

**Figure 2.**
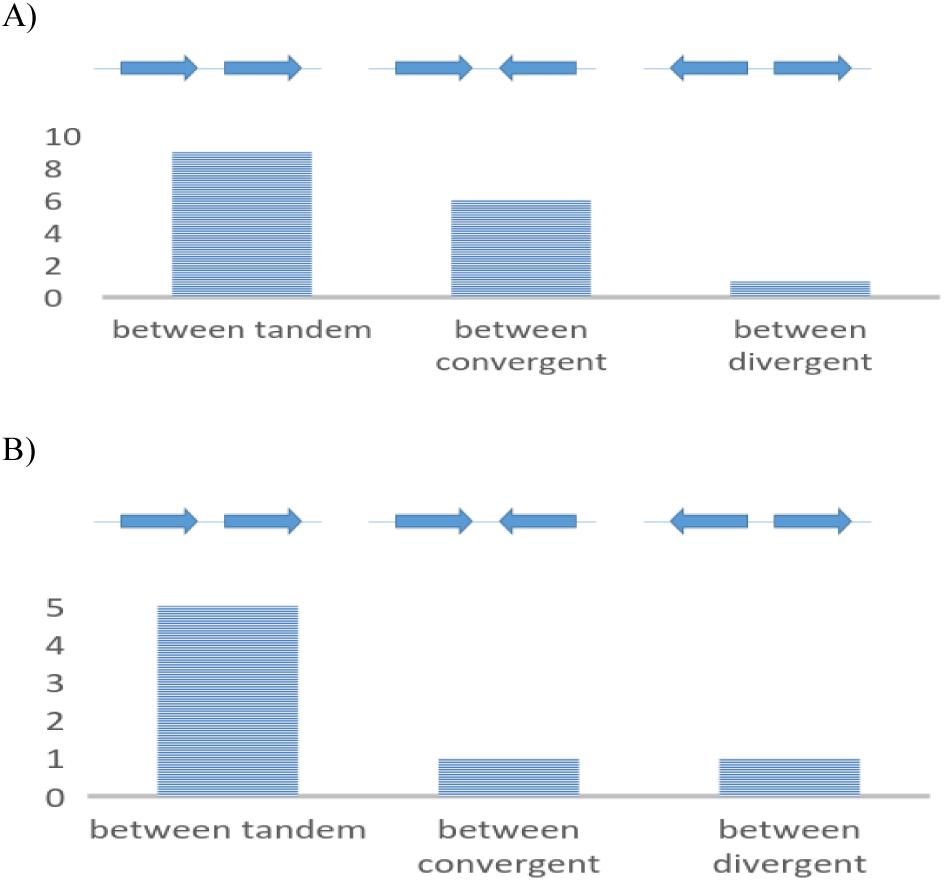
*Repeat counts of HPG1 (A) and HPG2 (B) members between tandem, convergent and divergent gene pairs*.

**Figure 3.**
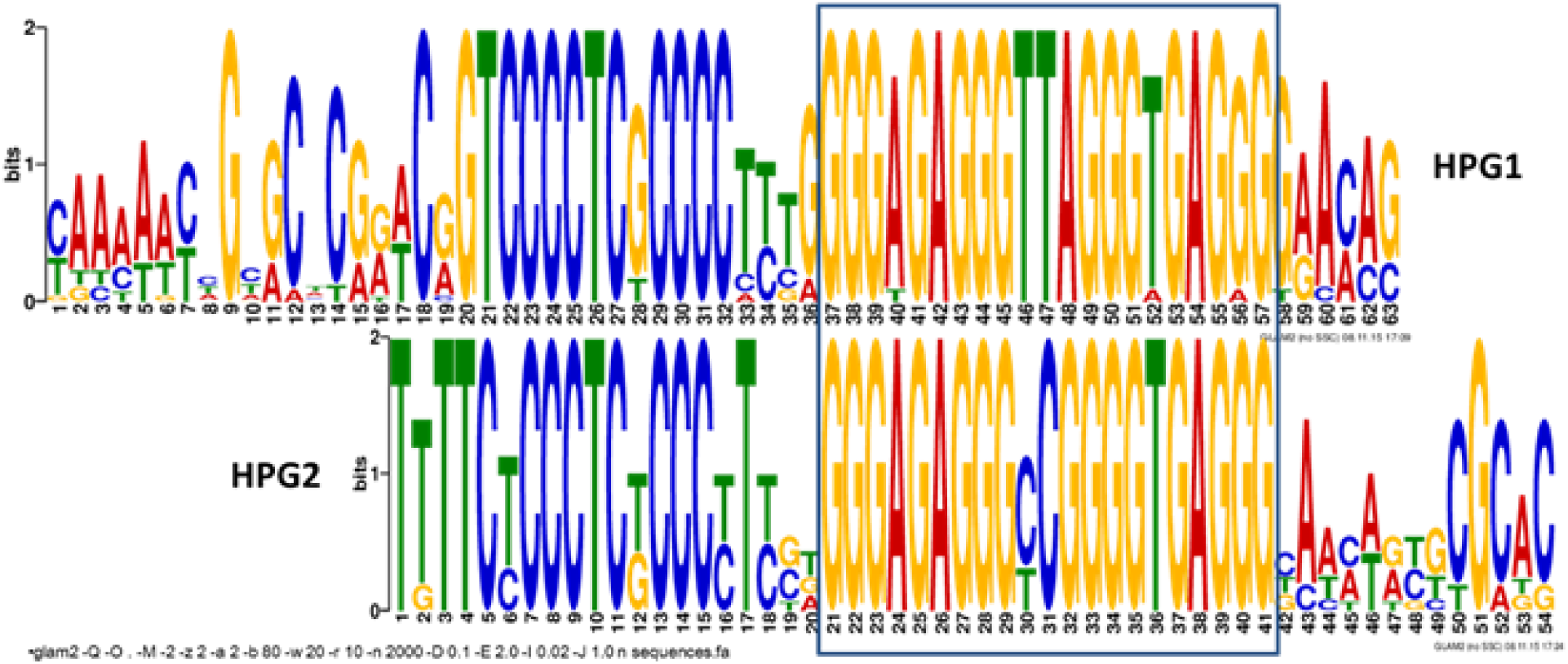
*Nucleotide allignment of HPGQ flanking regions for HPG1 (bottom) and HPG2 (top). GQ forming G-rich regions are alligned framed*.

The distances between ORFs are found to be between 377 nt and 65 nt. This small distances between ORFs complicates locating the putative regulatory regions between tandem gene pairs since most studies regard 200 nt upstream of the transcription start site as regulatory. Moreover, while the most of the HPGQs we identified are present between tandem gene pairs, 8 out of 16 HPG1s are found in the upstream of operons, indicating a role in the initiation of transcription.

For the HPGQs found between genes of the same operon a terminator role is plausible by blocking of RNA polymerase by GQs and prevention of further transcription. This is also supported by the previous studies indicating that polymerases “fall off” during transcription of GQ-induced DNA sequences (Siddiqui-Jain et al. 2002; Simonsson, Pecinka, and Kubista 1998) For HPG1, 6 of motifs found between convergent gene pairs while only a single is located between divergent gene pairs. The fact that HPGQs are found closer to the ORFs at the end may suggest this hypotheses further, however, it should be noted that previous studies have shown that non-B-motif formations may have regulatory effect even at a greater distance to the gene (Hatfield and Benham 2002).

The genomic distribution of HPGQs were also analysed according to each other. This revealed a surprising pattern. 6 of HPG1 and HPG2 members, HPGQs 5, 6, 32, 33, 34 and 35, were found to be located in close proximity (< 50 nt) and in pairs (Figure S2). Curiously these pairs were also located on the opposite strands. In addition, an HPG1 member, HPGQ 17 is also found in 50 nt upstream of HPGQ 18, a degenerate form of cHPG2. This indicates a similar function of HPG1 and HPG2. This is also supported by the sequence similarity of these two families.

### Search for conserved pattern in flanking regions

The close proximity of HPGQs arouses our suspicion for a conserved sequence pattern in the flanking regions. To reveal any conservation in the flanks we used GLAM2 tool from MEME suite (Bailey et al. 2009; Frith et al. 2008) to scan the sequences starting from 200 nt upstream to 200 nt downstream from the middle of the each HPGQ (401 nt in total). The results show that both of the HPGQ classes have a common C-rich feature at approximately 3 bases upstream of the HPGQs. The difference occur after the C-rich region – a highly conserved 4 nt long T-rich region precedes C-rich region for HPG2s. The alignment also reveals the similarity of HPG1 and HPG2 sequences – the difference of these groups peak at the composition of the second loop.

The close proximities of the members of HPG1 and HPG2s, as well as similar extended motifs is a strong indication that HPG1 and HPG2 are associated to same biological processes. For that reason, we decided to find a common motif for further analysis. Unfortunately, the motif output of GLAM2 tool is incompatible with other motif scanning and enrichment tools. This is because, unlike MEME, GLAM2 finds “gapped” motifs which reveals motifs even if there is insertion or deletions creating gaps in the motif. For that reason, a common motif was find for HPG1 and HPG2 using MEME tool (excluding a minority of the sequences that cannot be aligned properly due to “ungapped” alignment of the method). MEME analysis resulted in a 41 bp long motif with well-conserved C-rich and G-rich regions.

### Gene ontology (GO) analysis for the common motif

The motif obtained from MEME analysis using both HPG1 and HPG2 flanking sequences was used to find associated GO terms in *E.coli* using GOMo tool (Buske et al. 2010) by scanning between 1000 bp upstream and 200 bp downstream of first genes in operons. Among 208 predictions significant abundance in molecular function (MF) domain is observed. Three of top five predictions with %100 specificity was related to iron and sulfur cluster binding indicating a regulatory role of HPGQs (Table 2).

**Table 2.**
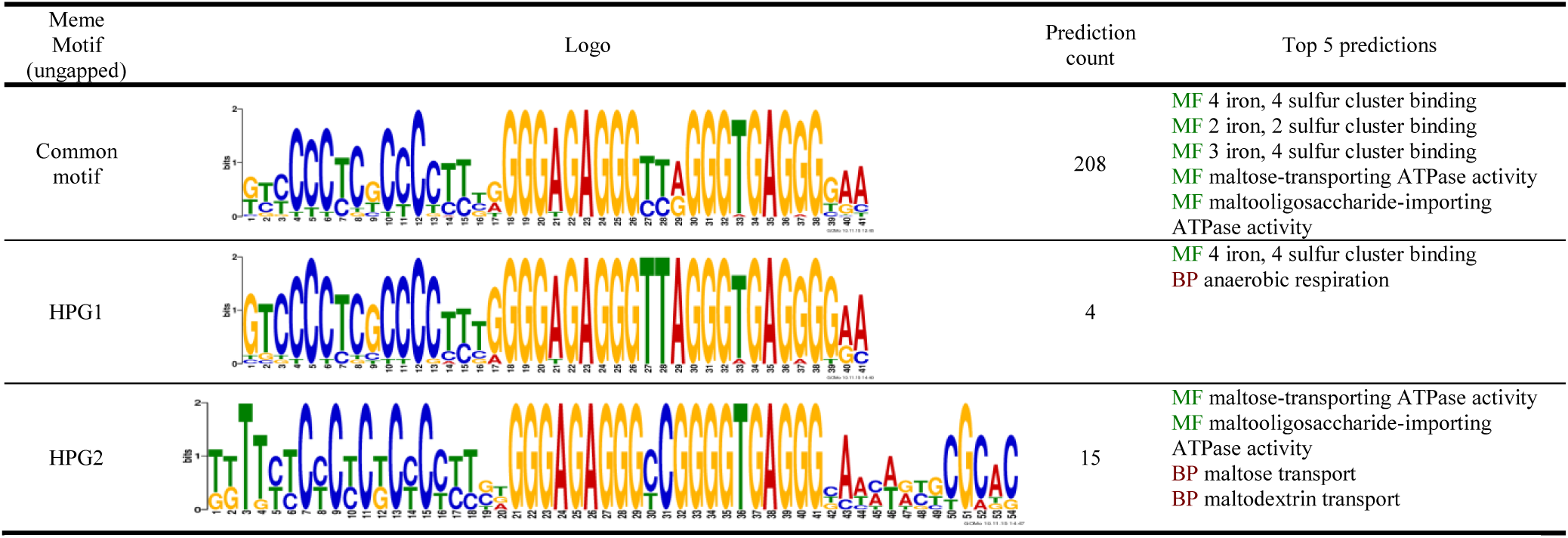
*GOMo analysis for prediction of associated GO terms linked to HPGQs*

| Meme Motif (ungapped) | Logo | Prediction count | Top 5 predictions |
| --- | --- | --- | --- |
| Common motif |  | 208 | MF 4 iron, 4 sulfur cluster binding<br>MF 2 iron, 2 sulfur cluster binding<br>MF 3 iron, 4 sulfur cluster binding<br>MF maltose-transporting ATPase activity<br>MF maltooligosaccharide-importing ATPase activity |
| HPG1 |  | 4 | MF 4 iron, 4 sulfur cluster binding<br>BP anaerobic respiration |
| HPG2 |  | 15 | MF maltose-transporting ATPase activity<br>MF maltooligosaccharide-importing ATPase activity<br>BP maltose transport<br>BP maltodextrin transport |

Interestingly, when HPG1 and HPG2 sequences were processed separately by MEME and the motifs found were used for the search of associated GO terms, HPG1-based motif was found to be associated with iron-sulfur cluster binding as well, while HPG2-based motif was found to be related to maltose transport, both being with 100% specificity. As a control, the motif was used to scan *Saccharomyces* genus, resulting in only a single prediction with specificity of 75% along with two predictions with negligible (0%) specificity.

It is important to note that higher discrepancy can be observed between motifs obtained by GLAM2 and MEME due to gaps present in C-rich region. This is especially noticeable for motif obtained from HPG2 sequences with their flanking regions.

### Search for degenerate HPGQs

G-quadruplex algorithm is capable of finding most of the G-quadruplex forming sequences however, the genome may also have degenerate forms of HPGQs mutated after loss of function. These sequences may not be able to be detected due to mutation at guanine sites. In order to reveal such sequences, the FIMO tool (Grant, Bailey, and Noble 2011) was used with the common motif in E.coli upstream sequences as well as alternate prokaryotic upstream sequences to determine a healthy the p-value cut-off. To our surprise, p-value plot (data not shown) revealed a significant increase beyond detected and additional three HPGQs. With a cut-off of 0.01, additional 3 motifs were found, indicating that motif scan may reveal additional but degenerate HPGQs (Table S2). It is also important to note that, the degenerate HPGQs may still form GQ with assistance of single guanines in the flanks or loops.

### HPGQ is not associated with any known motifs

The mode of mechanism of a regulatory motif may be dependent on transcription factor (TF)-binding ability of HPGQs in double stranded, hairpin or G-quadruplex state. It is expected that if the duplex formation is required for a TF to bind, for such a duplex-motif TF-binding would not depend on or require the preservation of a G-quadruplex formation and may only be an additional control mechanism to render the duplex-motif out of reach. In such a case we would expect to find such a motif preserved over G-quadruplex forming sequence. In other words, if such a motif is found, it can be suggested that the G-quadruplex formation may inhibit TF-binding rather than activation, given that the motif is preserved within G-rich region. We scanned for known motifs listed in CollecTF database (Kilic et al. 2014) within the common HPGQ motif. Using E-cut-off value of 0.1 the scan yielded no motifs and increasing the E-cut-off to 1 yielded only a single *E.coli* motif, binding site for Macrodomain Ter protein (MatP), with low probability, indicating that HPGQ motif is not associated with known motifs. However, it doesn’t mean a common TF may still be found in the vicinity that also has an undetected affinity for G-quadruplex or hairpin structures etc. Similarly no TF-binding motif is detected within 200 bp range of HPGQs by scanning for CollecTF motifs using Centrimo tool (Bailey and Machanick 2012).

### Topological variations of HPGQs

For a structural motif to have a direct influence on regulatory mechanisms, it is expected to be able to switch between its putative topologies, and evidently its level of interactions with intracellular factors such as TFs. In other words, for a form of regulation, structure should not be rigid or inert towards other factors and be able to switch between states. For GQs located on a duplex DNA, the B-form, in most cases, can be considered as an alternate state. In addition, the additional C-rich region conserved in the 5’ flank of HPGQs suggests formation of a hairpin together with two of the G-tracts. Thus, the HPGQ motifs may adopt the native B-form, G-quadruplex or hair-pin like structure. The plausibility of these states can be seen by comparison of free energies for each plausible topology. In order to find the nucleotides taking part in the GQ or hairpin structures and their free energies were calculated using ViennaRNA package (Lorenz et al. 2011) for both HPG1 and HPG2 motifs (Table 3). The free energies indicates close thermodynamic properties between hairpin and GQ structures indicating structural switch is possible. However, it should be remembered that the topologies are associated strongly with intracellular conditions, ligands, ions etc. Moreover, structural switch between alternate GQ topologies (*ie*. parallel vs anti-parallel) is also a possibility that may be revealed with further analysis.

**Table 3.**
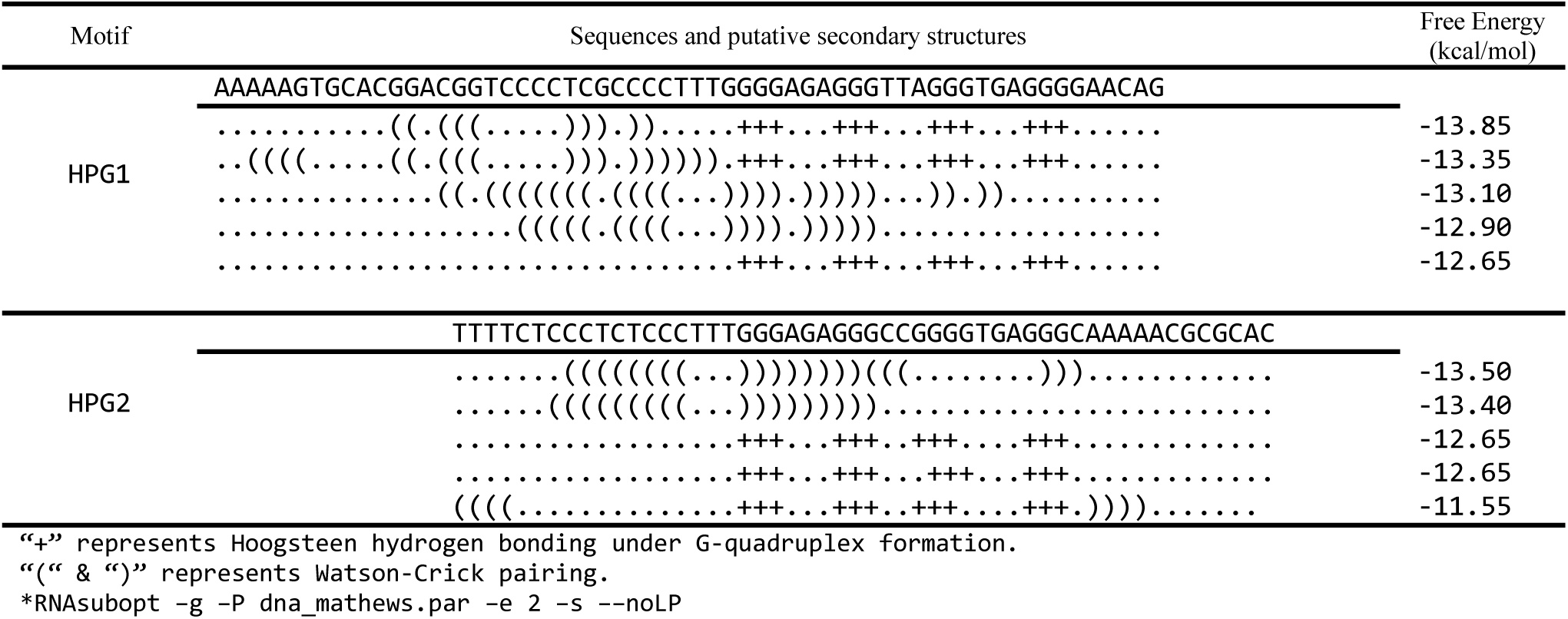
*Optimal and suboptimal secondary structures of common HPGQ motifs and the free energies of thermodynamic ensembles*.

## Discussion

Any intramolecular G-quadruplex forming sequence requires loop sequences between the G-tracts. Each following G-tract requires to fold on itself in intramolecular GQs and to align with other G-tracts. The necessary flexibility for the alignment is provided by this loop. Generally the loop requires to be longer than a single nucleotide and usually no longer than seven. Because the role of the loop is mainly steric, it is expected that the sequence of the loops are less effective than the stem on the formation of GQs. Indeed, most studies show that the loop composition has marginal effect on the stability of the structures, while the length is highly influential on topology as well as stability. However, when considering a biological function of a GQ, we need to consider the whole topology, not only the ability to form a G-quadruplex. In this case, the loop compositions may have direct role by interacting other biological molecules. For that reason we hypothesized that if a G-quadruplex have a direct interaction with proteins, the loop sequences should be conserved between motifs of similar origin or function.

With this aim, for the first time, we have applied a classification for putative G-quadruplex forming sequences based on phylogenetic analysis to put weight on the loop sequences when studying GQs. Our analysis on *E.coli* genome revealed two groups of putative G-quadruplex forming motifs by scanning for putative GQs using a generally accepted algorithm which allows freedom to the loop sequence. Sequences are then grouped using a phylogenetic clustering web tool revealing two major groups with very high sequence similarity. Interestingly these two groups consisting almost half of the putative GQs found and are located between prokaryotic open reading frames and most of them are between operons indicating a regulatory role. Since the loop sequence influences the final topology of the G-quadruplex and its ability to form H-bonds with other biomolecules, the strong conservation of loops clearly indicates that biological significance of the G quadruplexes is not only comes from the formation of G-tetrad but the formation of a specific G-quadruplex topology. For that reason, investigation of any putative G-quadruplexes without classification overlooks the potentially different roles of G quadruplexes.

Genomic distribution analysis of these sequences showed strong correlation with each other, since they are often found in pairs and on opposite strands. This was also supported by sequence alignments when all members of both HPGQ groups shared a C-rich region in the 5’ flank, as well as strong similarity in G-rich region except at the middle loop position. The conserved C-rich region also revealed possible formation of hairpin structures. This indicates that HPGQs may be able to switch, not only between GQ and duplex DNA but also, a hairpin-like topology.

It should be remembered that the G-quadruplex algorithm did not allow any flexibility on the unity of the guanine-tracts, in other words, any sequences similar to HPGQ except an insertion/deletion mutation inside the guanine tract would not be found. In order to include these we have scanned the whole genome using the motif revealing three additional degenerate HPGQs. It is important to note that these sequences could fit our algorithm with a single insertion/deletion (indel) mutation in the G-rich region indicating a functional role of G-quadruplex structure. Moreover, it is not possible to eliminate G-quadruplex formation for degenerate forms since it is known that singled-out guanines in the flanks may take part formation of G-tetrads.

The aligning all HPGQs together with their flanking sequences led to a common motif. The presence of a conserved C-rich region at the 5’ flank rose our suspicion for the formation of non-GQ topologies, particularly hairpin formation. Thermodynamic calculations showed that both GQ and hairpin structures are possible to form and may switch between two topologies. Such a transition is an indication of an active role in the regulatory mechanism. Similar transitions between GQ and hairpins have been shown to have biological relevance previously (Kuo et al. 2015; Romanucci et al. 2015). However, further laboratory work is required to clarify potential topologies as well as the regulatory mechanism of HPGQs found in this study.

## Methods

The genome of E.coli K12 MG1655 was obtained from NCBI database and searched for putative G-quadruplexes using quadparser software (Wong et al. 2010) and the standard pattern G*_S_*N*_L1_*G*_S_*N*_L2_*G*_S_*N*_L2_*G*_S_* where G refers to guanine and N refers to any nucleotide including guanines. While s is set to be equal to or bigger than 3, and L1-3 are smaller than 7 but bigger than 0. The program was run using both + and − strands. The sequences and their starting positions were listed and confirmed using NCBI sequence viewer (Aiyar 2000; Higgins and Sharp 1988; Sievers and Higgins 2014; Wolfsberg 2011). Then the sequence list was processed using Clustal Phylogeny web tool (Larkin et al. 2007) for sequence alignment and phylogenetic tree generation using UPGMA as the clustering method instead of default neighbour-joining method (Saitou and Nei 1987; Zhang and Sun 2008). We allowed a single indel mutation inside the minimal G-quadruplex forming sequence for each group members. Any HPGQ with similar sequence with less than two indel mutation was considered to be members of same group. This corresponded to the phylogenetic distance threshold of 0.047, the ration of the number of substitutions per the length of HGPQ. Non-clustered HPGQs were not taken into consideration for following studies. The locations of the two main HPGQ groups were mapped according to the gene ORFs on the genome as defined on NCBI database. The distance to the genes and their directions were listed. The order of the genes in the operon was taken from (Salgado et al. 2013).

### Motif Discovery and Enrichment

Selected HPGQ sequences and their flanking regions (±200 bp) were used to motif finding studies. In order to reveal other conserved motif in the flanking regions we used GLAM2 web tool (Frith et al. 2008). We chose GLAM2 over MEME because such a motif do not have to have the identical distance to the G-quadruplex forming sequence and GLAM2 is capable of finding “gapped” motifs. HPG1 and HPG2 are scanned separately using only the G-quadruplex forming strand and the maximum number of columns to be aligned was set to 80, instead of default value, 50. Because GLAM2 returns a non-compatible motif format to be used for motif enrichment tools such as GOMo, we also applied HPG1 and HPG2 to MEME motif finder tool, both separately and combined (common motif) using the same parameters as mentioned for GLAM2. Discovered motifs from were scanned for associated gene ontology terms using GOMo using default parameters (Buske et al. 2010). In order to discover any degenerate HPGQs with similar sequence but not detected due to restriction of G-quadruplex algorithm, we scanned the *E.coli* K12 MG1655 uid57779 upstream sequences for the common motif using FIMO web tool. Centrimo was used to discover associated motifs and transcription factor binding sites in the flanking regions. The HPGQ sequences and their flanking regions (±200 bp) were scanned for motifs from the CollecTF (bacterial TF motifs) database using default options(Kilic et al. 2014).

### Secondary Structure Search

Secondary structural modelling was performed using ViennaRNA package (Lorenz et al. 2011). Initially, two sequences were obtained to represent HPG1 motif and HPG2 motif. This was done using RNAalifold (Bernhart et al. 2008) and the sequence alignment by GLAM2 yielding two sequences with 63 and 54 bases long, respectively. Next, the sequences were scanned for hairpin and G-quadruplex formation by RNAsubopt using “no lonely pair” (--noLP), and “scan for g-quadruplex” (-g) parameters. It is important to note that the default parameter file is for RNA secondary structures. For that reason we specified the parameter file obtained from RNAalifold Webserver modified for DNA structures (Gruber, Bernhart, and Lorenz 2015; Mathews 2004). Chosen secondary structures and their free energies were listed to represent various topologies.

